# Nectins rather than E-cadherin anchor the actin belts at cell-cell junctions of epithelia

**DOI:** 10.1101/809343

**Authors:** Pierre Mangeol, Dominique Massey-Harroche, André Le Bivic, Pierre-François Lenne

## Abstract

Cell-cell junctions support the mechanical integrity of epithelia by enabling adhesion and tension transmission between neighboring cells. The prevailing mechanistic dogma is that E-cadherin supports and transmits mechanical tension between cells through actin belts in a region named the *zonula adherens*. Using super-resolution microscopy on human intestinal biopsies and Caco-2 cells, we show that the *zonula adherens* consists of E-cadherin and nectin belts that are separated by about 150 nm along the apico-basal direction, the nectin belt being in the immediate vicinity of the actin belt. The segregation of nectins and E-cadherin increases as the tissue matures. Our data redefine the structure of the *zonula adherens* and show that nectins, rather than E-cadherin, are the major connectors of actin belts in epithelia.

## Main text

The current description of epithelial apical junctions (*1*) is based on the seminal work of Farquhar and Palade using electron microscopy (*2*), which defined three different areas in apical junctions of enterocytes: the most apical *zonula occludens*, now known as tight junctions, followed basally by the *zonula adherens* (*ZA*), with a wider intercellular space and the accumulation of cytoskeletal fibers, and the most basal *macula adherens*, consisting of desmosomes (*3*). Molecular constituents were identified later, such as E-cadherin (E-cad) (*4*, *5*), that was found on the lateral membrane, with an accumulation at the *ZA* (*6*). Identification of the E-cad/β-catenin/α-catenin complex followed shortly after (*7*), and because of α-catenin interaction with actin (*7*, *8*), the E-cad-catenin complex entered textbooks in the early 90s as the complex linking the actin meshwork of neighboring cells (*9*).

However, another major protein complex including a transmembrane protein and an actin binding protein was discovered a decade later, the nectin-afadin complex (*10*, *11*). This complex appears to be more specific to the *ZA* than the E-cad-catenin complex (*11*). Afadin has been shown to be essential during development, including during the mechanically demanding phase of gastrulation, both in mouse (*12*) and *Drosophila* (*13*); afadin is also critical for epithelial apical constriction (*13*). Similarly, nectin-2 was found to be crucial during epithelial apical constriction of the neural tube in *Xenopus*: nectin-2 depletion inhibits the epithelial apical constriction, whereas its over-expression enhances it (*14*). Together, this shows the functional importance of nectins and afadin for epithelia during organ development.

In the last years, the classical view of E-cad-catenin complex interaction with actin has been challenged. In mammary cells, actin organizes around E-cad clusters rather than clearly connecting to them (*15*), and *in vitro* experiments showed that E-cad-catenin binds actin efficiently only under tension (*16*). To date, both E-cad and nectin-based protein complexes are thought to be located at the *ZA*, and both have the potential to physically connect the actin meshwork from one cell to its neighbors, but their respective roles have not been clearly discriminated.

To obtain a detailed view of this essential structure, we decided to use super-resolution microscopy and chose to investigate the supramolecular organization of the *zonula adherens* in human intestinal biopsies and human intestine enterocyte-like Caco-2 cells (*17*). We imaged these samples with Stimulated-Emission-Depletion (STED) microscopy (*18*), improving the spatial resolution by 3-fold in the junctional plane and 7-fold along the apico-basal axis compared to classical confocal microscopy approaches.

To get a better understanding of the organization of the *ZA* in intestinal cells, we started by imaging E-cad and filamentous actin (F-actin) in Caco-2 cells. Cells were seeded on filters and grown over 14 days to allow differentiation (*19*). Similarly to what has been observed before (*6*, *20*, *21*), we found F-actin and E-cad on the lateral side of cells, and both accumulate at the most apical part (Fig. 1A-B). The accumulation of F-actin corresponds to the actin belt that can be readily observed when imaging cells from their apical side and it can be well distinguished from the terminal web (Fig. S1A). When focusing on the apical region with STED, we found that E-cad accumulates about 100 to 300 nm basally from the actin belt (Fig. 1A-B). E-cad is in the vicinity of actin structures, but these actin structures have an intensity about 6 times weaker than the intensity of the actin belts when measuring along the apico-basal axis (Fig. S1B). These results are surprising because according to the classical model of the *ZA*, E-cad and catenins tie together the actin belts of neighboring cells and should therefore localize at the same level. To confirm this result, we imaged both α-catenin and β-catenin in Caco-2 cells and made the same observation as for E-cad: the entire E-cad-catenin complex is located about 100 nm basally from the actin belt (Fig. 1A,B).

**Figure 1.**
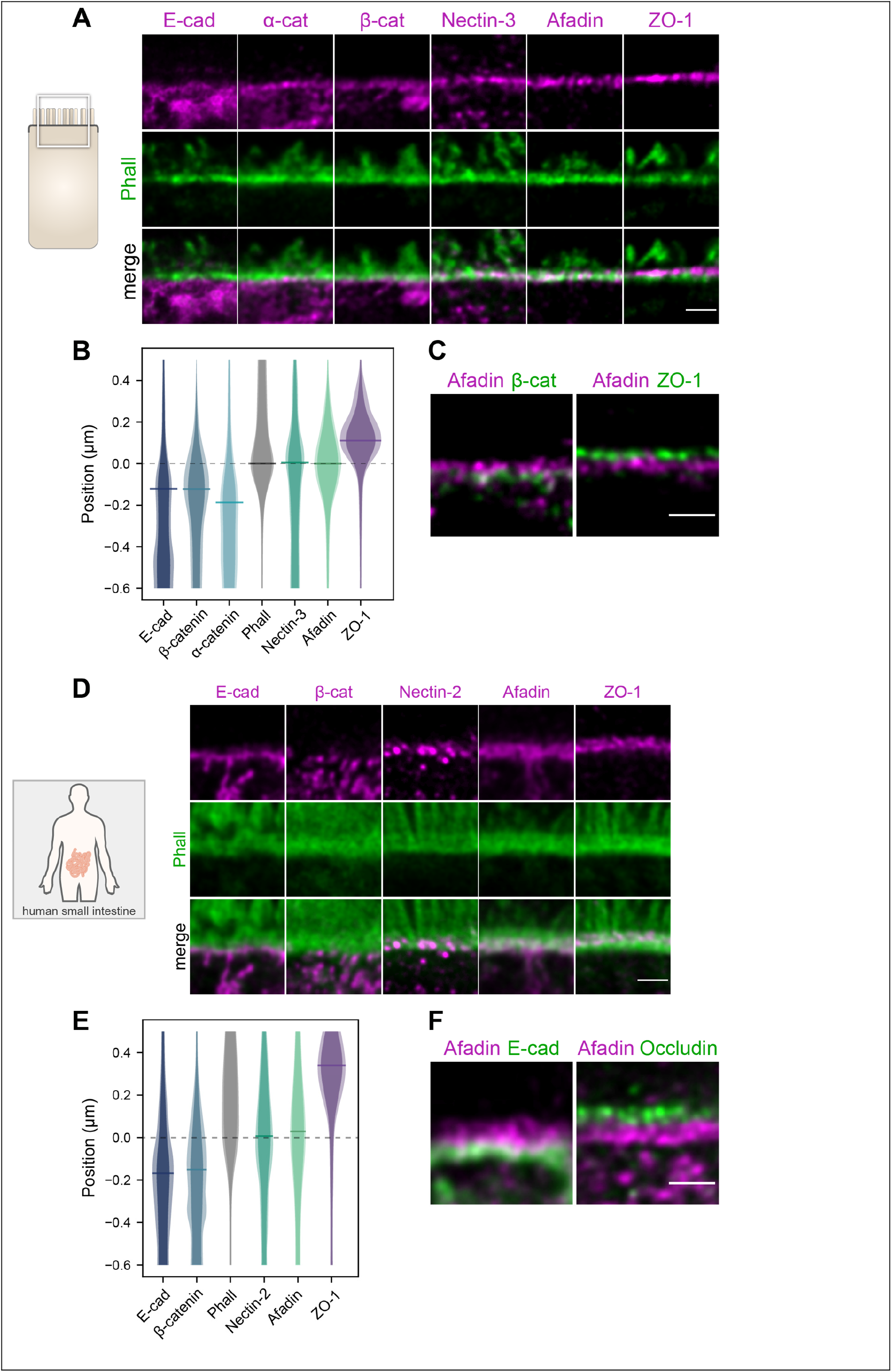
Segregation of adhesive complexes and their relative position compared to the actin belt in Caco-2 cells (A-C) and human ileum biopsies (D-F). **(A,D)** Side view images of junctional complexes proteins (magenta) and F-actin (green). **(B,E)** Quantification of the position of adhesive complexes with respect to the actin belt on the apico-basal axis. The distributions represent the average intensity of a given protein labelling (plain color) and the average added with their standard deviation (lighter color). Bars on distribution indicate the maxima of the distributions. Methodological details are given in the Supplementary Materials. **(C)** Segregation of the tight junction (ZO-1), afadin and β-catenin in Caco-2 cells. **(F)** Segegation of the tight junction (occludin), afadin and E-cad in human ileum biopsies. Scale bars 1 μm.

These unanticipated results led us to wonder whether nectins rather than E-cad could connect actin belts in adjacent cells. When observing nectin-3 and afadin in Caco-2 cells, we found that both proteins are precisely located at the position of the actin belt on the apico-basal axis (Fig. 1A,B). Consistent with these results, we found that afadin and β-catenin are present in distinct regions, seemingly organized as belts, where afadin is apical of β-catenin (Fig. 1C). The tight junction is present further apically, as ZO-1 forms a distinct belt located apically of afadin (Fig.1A-C). We also detected nectin-1 and nectin-2 in the same region (Fig. S1C). Altogether, these results show that two distinct belts of proteins define the *ZA*: the afadin and nectins belt that align with the actin belt along the apico-basal axis, and the more basal E-cad and catenins belt. Each belt has a similar width of about 100 nm on average along the apico-basal axis. The tight junction and the E-cad-catenins belts are not devoid of filamentous actin, which is, however, significantly scarcer than at the nectin-afadin belt. The adhesive complex transmitting force between actin belts of neighboring cells has to align with these belts, because of mechanical balance. Therefore, our results suggest that the nectin-afadin complex is responsible for tension transmission at the *ZA* rather than the E-cad-catenin complex.

This unexpected segregation of E-cad and nectin complexes in human intestinal cells in culture prompted us to investigate how E-cad and nectins are organized *in vivo*. To uncover this organization, we observed the localization of E-cad-based and nectin-based protein complexes in human small intestine (ileum) biopsies. We started by observing the overall localization of these proteins. We found E-cad and catenins at the junctions of all epithelial cells (Fig. S2A), whereas nectins and afadin showed a different pattern of localization. While nectin-2 and afadin labelled all epithelial apical junctions, nectin-1 and nectin-3 labelled the apical junctions of cells in crypts (Fig. S2B), similarly to what has been reported in the Human Protein Atlas (*22*). The organization of actin was also drastically different between villi cells in which F-actin accumulates at the apical junction and crypt cells in which such an accumulation is not obvious (Fig. S2C). For simplicity, we limited our study to enterocytes in villi. In these cells, we observed that along the apico-basal axis, F-actin is often organized as a thick belt, of about 300 nm in width (Fig. 1D). When we observed the organization of the *ZA* proteins, similarly to what we observed earlier in Caco-2 cells, E-cad-catenin complexes were separated from nectin-afadin complexes (Fig. 1D-F) and localized more basally. Here, nectin-2 and afadin align with the lower part of the actin belt along the apico-basal axis. In contrast, the tight junction is localized in the higher part of the actin belt (Fig. 1D-F).

Earlier studies have reported that nectins and afadin localize in the same region as E-cad (*10*, *11*) or that nectins are localized at the tight junction (*23*). The discrepancy between our findings and earlier studies may stem from two non-exclusive possibilities: the lower spatial resolution of the previously used techniques or the maturation of the cells. These studies often used cell cultures that had just reached full confluence, a state obtained a few days after seeding cells on a substrate. At this differentiation stage cells have junctions, but these often do not recapitulate the functional characteristics found *in vivo*, which usually requires more time (*19*). To test whether this co-localization is due to a lack of resolution or to a potential maturation of the junctions during the epithelial layer maturation, we seeded Caco-2 cells on filters and fixed them at intermediate stages of maturation: just after they reached full confluence at 6 days and later at 9 days. In comparison, the results shown on Fig.1 are obtained from cells at 14 days (*i.e.* being fully confluent for 8 days). Cells that had just reached full confluence, at 6 days, showed a much lower segregation of E-cad and afadin compared to more mature cells (Fig. 2A,B). The tight junctions were also in close vicinity to the adhesive complexes, but could already be discerned (Fig. 2A,B). At 9 days, the segregation of E-cad and afadin was more pronounced, although it was still less than at 14 days (Fig. 2C,D). At 9 days, a clear gap between E-cad and the tight junction could be observed (Fig. 2C). Afadin and β-catenin were often difficult to separate at 6 days, whereas at 9 days they appeared partly segregated (Fig. 2E). Taking these results together, we found that as cells mature, the tight junction, the nectin-based and the E-cad-based junctions segregate, and F-actin accumulates at the level of the nectin-based junction (Fig. 2F).

**Figure 2.**
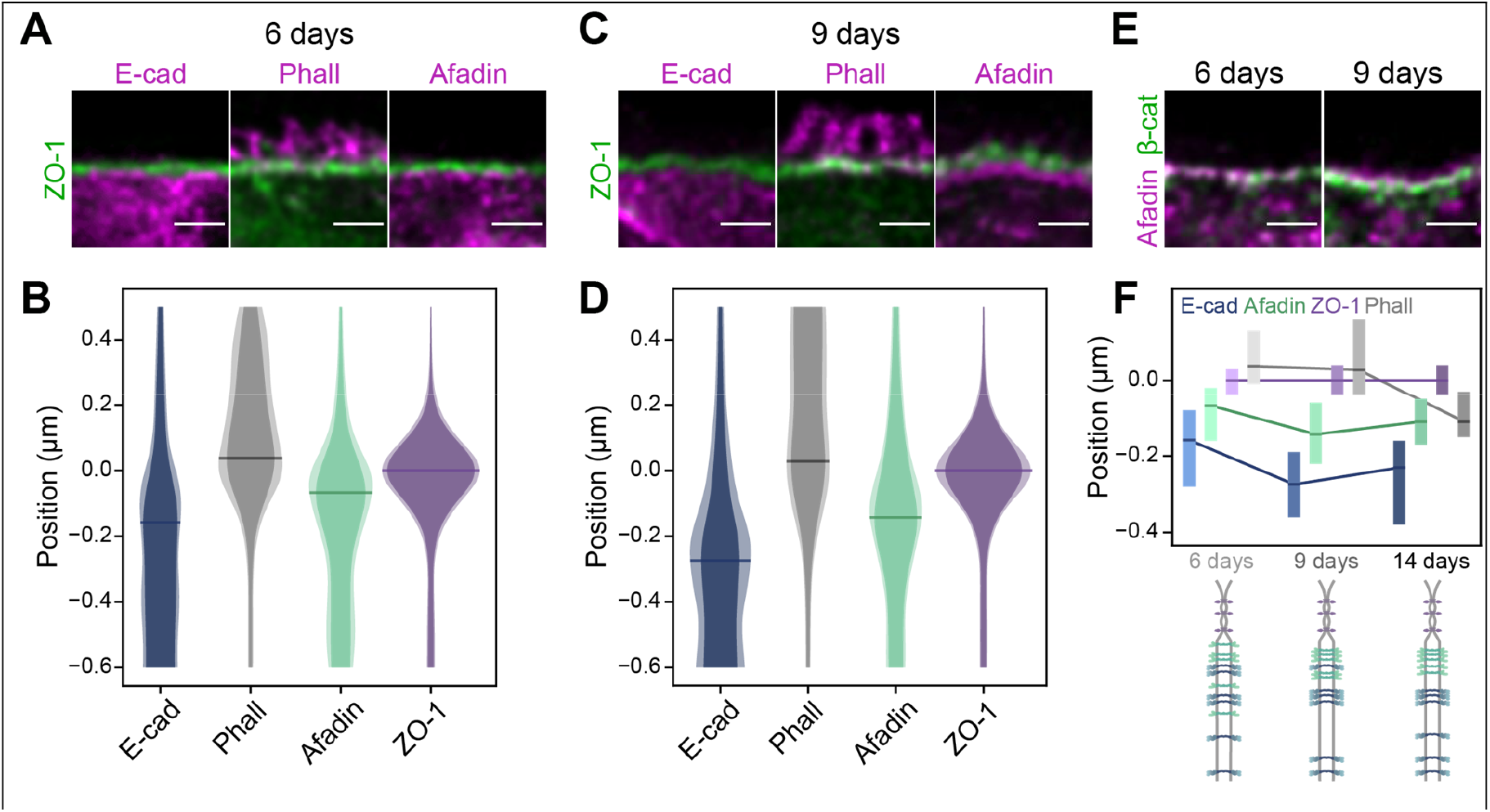
The segregation of the adhesive complexes intensifies as cells mature. (**A**,**C**) Side view images of Caco-2 cells apical junctions comparing the tight junction (ZO-1) position to the position of E-cad, F-actin and afadin at 6 days (A) and 9 days (C). At these stages most of F-actin signal is concentrated in microvilli. (**B**,**D**) Quantification of the position of the previous proteins with respect to ZO-1 on the apico-basal axis at 6 days (B) and 9 days (D). Representations of distribution identical to Fig. 1. (**E**) Afadin (magenta) and β-catenin (green) segregate as cells mature. (**F**) The segregation of adhesive complexes intensifies over several days after cells reached full confluence. Top, quantification of protein localization; bars represent the 30% most frequent localization of a given protein at +/− 0.5 µm the position of center of the tight junction along the apico-basal axis. Bottom, schematic of the adhesive complexes localization in the apico-basal direction; tight junction in magenta, nectin-afadin in green, E-cad-catenins in blue. Scale bars 1 µm.

How can maturation influence the segregation of adhesive complexes? In cell culture, when two cells contact each other and initiate junction formation, *i.e.* on short timescales, E-cad and nectins are both present at the nascent junction (*24*), presumably competing to interact with actin via their adaptor proteins. However, what happens after several days remains unknown. The E-cad-catenin interaction with actin strongly depends on the force applied to the bond *in vitro* and this bond has at best a lifetime in the order of a second or less in a narrow force regime around 8 pN (*16*). This suggests that the interaction may not be stable in the long term, in particular if the E-cad-catenin complex has to compete with other complexes, such as nectin-afadin. Although isolated afadin and α-catenin have a similar affinity constant for F-actin (*8*, *10*), the kinetics or the force dependence of the nectin-afadin interaction with actin remains to be determined. Additionally, collective processes such as clustering of adhesive complexes could reinforce the segregation of nectins from E-cad (*25*). Our data suggests that on the timescale of days the nectin-afadin complex could be better suited to link the actin belts of neighboring cells than the E-cad-catenin complex.

We wondered if we could observe the structural link between actin belts and the nectin-afadin complex. To this end, we carefully observed the junction area in the planar orientation. Actin belts of neighboring cells can be individualized with STED microscopy (Fig. 3A,B). The distance between actin belts of adjacent cells is in the range of 250-400 nm. When observing further these images, we found actin connectors extending from actin belts towards the neighboring cell membrane, in the form of facing combs (Fig. 3A,B). Remarkably, we frequently found afadin organized as puncta in the immediate vicinity of these F-actin connectors, seemingly linking actin connectors between neighboring cells (Fig. 3B,C). These results suggest that afadin together with nectins link neighboring cells actin belts using F-actin connectors.

**Figure 3.**
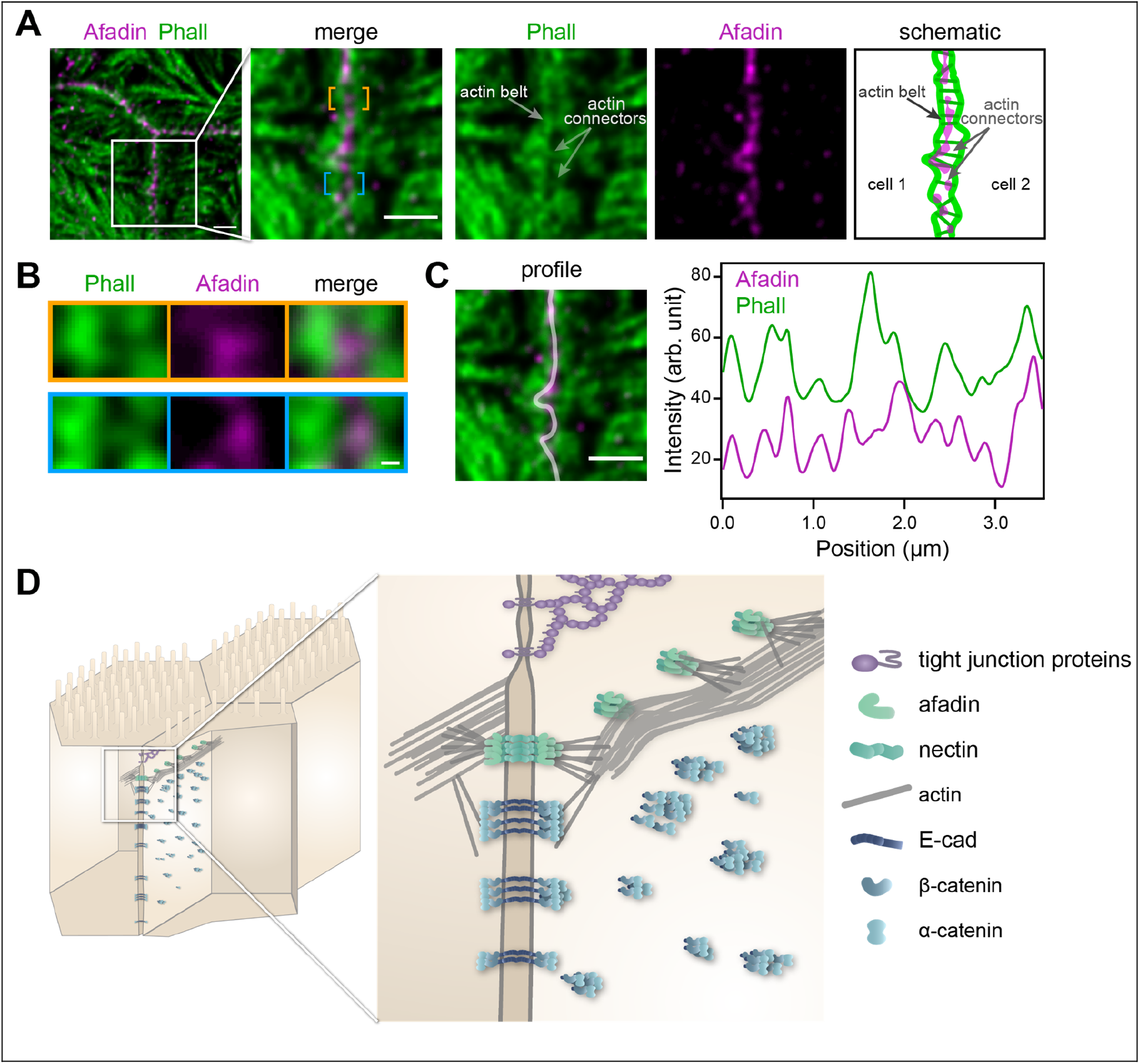
Afadin is present on actin structures connecting actin belts. **(A)** Planar view of cells junctions labelled with phalloidin (green) and anti-afadin (magenta). Actin belts, actin connectors (in the vicinity of afadin staining) as well microvilli can be observed. Right, schematics of the labelling, in which actin belts, actin connectors and afadin staining are represented. **(B)** Zoom in the regions of interest indicated by a yellow and blue brackets on the merged image in (A). Actin connectors as well as afadin puncta can be observed. **(C)** Profile on the junction (left) and corresponding intensity profile along the junction (right). The profile is taken from bottom to top, following afadin labelling, showing a fair coincidence between the afadin and F-actin signals. **(D)** Model of the *ZA* of mature junctions. The organization of actin together with afadin may take several forms, *i.e.* direct binding of afadin to the actin belt, or through actin connectors. (A,C) scale bars 1 μm. (B) scale bar 100 nm.

Altogether, our results show that the classical model of the *ZA* must be redefined. First, we found using STED microscopy both *in vivo* and in cell culture that the nectin-afadin complex is precisely located at the position of the actin belt on the apico-basal axis in intestinal epithelial cells. In contrast, the E-cad-catenin complex is found more basally without obvious physical link to the apical belt. Second, our cell culture time course demonstrated that the organization of the *ZA* depends on cell maturation, as the segregation between nectin-afadin and E-cad-catenin intensifies over 14 days. In immature junctions, it is likely that both E-cad and nectin adhesive complexes are responsible for linking neighboring cells actin belts. Third, our results showed that the nectin-afadin complex is concentrated precisely where actin belts are linked, whereas the E-cad-catenin complex is mostly excluded from these regions. Therefore, we propose a new model for the *ZA* where in immature junctions both the nectin-afadin and E-cad-catenin complexes link neighboring cells actin belts, whereas in mature junctions the nectin-afadin complex is responsible for linking actin belts of neighboring cells (Fig. 3D). The precise organization of F-actin together with afadin is unknown, but the fact that afadin binds F-actin to its side rather to its end (*10*) is likely to play an important role in this organization. This redefinition of the *ZA* will help to better understand why nectins and afadin have crucial roles when epithelia are mechanically challenged (*12*–*14*). Moreover, it redefines the role of the E-cad-catenin complex that appears to be responsible for the overall cell-cell adhesion but may have a secondary role in transmitting tension between epithelial cells, at least when nectins are present. A possible explanation of our results could be the following: when a junction is established, because the E-cad-catenin complex binds F-actin efficiently only under tension (*16*), its association with actin belts may be only transient, whereas other protein complexes that have a more stable interaction with actin could take over as the junction matures. Our study that highlights an unanticipated structural role of nectins and afadin suggests new avenues to explore their role in the dynamics of cellular adhesion, and could benefit to the mechanistic understanding of nectins implication to multiple human pathologies (*26*).

## Acknowledgment

We would like to thank Marc Lopez for the gift of several antibodies, Flora Poizat for human biopsies, Fabrice Richard, Virgile Viasnoff, Stefan Harmansa, Emily Gehrels, Anaïs Bailles, Frank Schnorrer and his group, Le Bivic and Lenne groups for discussion. Biopsies were obtained with the agreement IPC-CNRS-AMU 154736/MB.

## Funding

We acknowledge the IBDM imaging facility, member of the national infrastructure France-BioImaging supported by the French National Research Agency (ANR-10-INBS-04). PM was supported by ITMO Cancer (Plan Cancer), Ligue nationale contre le Cancer and the French National Research Agency (ANR-T-JUST, ANR-17-CE14-0032). The project developed in the context of the LabEx INFORM (ANR-11-LABX-0054) and of the A*MIDEX project (ANR-11-IDEX-0001-02), funded by the “Investissements d’Avenir” French Government program

## Author contributions

P.M. designed and performed the experiments, and analyzed the data. D.M-H. assisted in performing the experiments. P.M., A.L.B and P-F.L. acquired financial support for the project. All authors discussed the results and contributed to the manuscript

## Competing interests

Authors declare no competing interests.

