## Supplementary Materials for "Nectins rather than E-cadherin anchor the actin belts at cell-cell junctions of epithelia"

### Methods

#### Cell culture

A clone of Caco-2 cells, TC7, was used in this study because differentiated TC7 cells form a regular monolayer (*1*). Cells were seeded at a low concentration of $10^5$ cells on a 24 mm polyester filter with 0.4 µm pores (3450, Corning inc., Corning, NY). Cells were maintained in Dulbecco's modified Eagle's minimum essential medium supplemented with 20% heat-inactivated fetal bovine serum and 1% non-essential amino acids (Gibco, Waltham, MA), and cultured in 10% $CO_2$/90% air. The medium was changed every 48 hours.

#### Sample preparation for immunostaining

##### Human sample preparation

Human intestine biopsies were obtained under the agreement IPC-CNRS-AMU 154736/MB. Intestinal samples were fixed in paraformaldehyde (PFA 32%, Fischer Scientific) 4% in phosphate buffer saline (PBS, Gibco) for 4 hours at 4°C. Biopsies were then cryoprotected with sucrose (10% sucrose in PBS for 45 minutes, 20% sucrose for 45 minutes followed by 30% sucrose overnight at 4°C). Biopsies were finally embedded in optimal cutting temperature compound (OCT compound, VWR) and frozen in liquid nitrogen.

##### Cell culture preparation for optical microscopy

Cells were washed in PBS and then fixed in PFA 4% in PBS for 20 minutes at room temperature. When apico-basal orientation observations were needed, cells were sectioned along the apico-basal axis. Prior sectioning, cells were embedded in OCT compound and frozen in liquid nitrogen.

##### Samples sectioning

All samples were sectioned with a cryostat (Leica CM 3050 S, Leica Biosystems). 10 µm sections were transferred to Superfrost Plus adhesive slides (Thermo Scientific) prior labelling.

##### Immunostaining

Intestinal sections and cultured cells were prepared similarly. Samples were permeabilized in 1% Triton X100 (Sigma-Aldrich) in PBS for 10 minutes. After washing with PBS, samples were saturated with 10% fetal bovine sera (Gibco) in PBS (“saturation buffer”) over an hour at room temperature. Primary antibodies were diluted in the saturation buffer and incubated overnight at 4°C. In more details: rabbit anti-ZO-1 (1/500, 61-7300, Invitrogen), mouse anti-occludin (1/500, 331500, Invitrogen), mouse anti-E-cadherin (1/500, 610181, BD Biosciences, used on Caco-2 cells), rat anti-E-cadherin (1/400, M108, Takara, used on human biopsies), rabbit anti-β-catenin (1/200, ab32572, Abcam), rabbit anti-alpha-catenin (1/200, 71-1200, Invitrogen), mouse anti-afadin (1/100, 610732, BD Biosciences, used on Caco-2 cells), rabbit anti-afadin (1/200, HPA030213, Sigma-Aldrich, used on human biopsies), mouse anti-nectin-1 and anti-nectin-3 (1/200, kind gifts of Marc Lopez), rabbit anti-nectin-2 (1/200, HPA012759, Sigma-Aldrich). Secondary antibodies were incubated 1 hour at room temperature. Alexa Fluor 568 conjugated to antibodies raised against mouse, rabbit and rat and Alexa Fluor 532 conjugated to antibodies raised against mouse and rabbit (Invitrogen) were used at 1/200 dilution in the saturation media. Phalloidin Alexa Fluor A532 (Invitrogen) was mixed with secondary antibodies and used at 1/100 dilution. After each incubation, samples were rinsed 4 times with PBS. Samples were finally mounted in Prolong Gold antifade mountant (Invitrogen) at 37°C for 45 minutes.

#### STED microscopy

Images of samples were acquired with a STED microscope (Leica TCS SP8 STED, Leica Microsystems GmbH, Wetzlar, Germany), using a 100X oil immersion objective (STED WHITE, HC PL APO 100x/1.40, same supplier). Two-color STED was performed with Alexa Fluor 532 excited at 522 nm (fluorescence detection in the 532-555 nm window), and Alexa Fluor 568 excited at 585 nm (fluorescence detection in the 595-646 nm window). To minimize the effect of drifts on imaging, both dyes were imaged sequentially on each line of an image and depleted using the same 660 nm laser. Detection was gated to improve STED signal specificity.

#### Analysis of protein density along the apico-basal axis (Fig 1B,E, 2B,D)

In order to quantify the density of proteins along the apico-basal axis and their positions with respect to each other, we used custom-made Image-J macros and Python programs. In brief, in each case a reference protein was localized precisely, defining a reference position along the junction from which intensity measurement was done. The intensity of protein labelling was measured along the apico-basal axis for each position of the junction. In practice, every 10 nm along the junction, an algorithm drew a segment along the apico-basal axis to extract the intensity profile of both proteins studied. In the process, we used bilinear interpolation to obtain sub-pixel quantification. Results of analyses were then normalized for each junction to avoid junction-to-junction intensity variation. Because we used a reference protein for each junction, we could then align all results based on the reference position of the reference protein and pool all results into a single protein density plot.

For Figure 1B (Caco-2 cells 14 days) the reference was the actin belt, which position was found automatically with an ImageJ macro, determining the maximum intensity of Phalloidin staining intensity along the apico-basal direction for each position along the junction.

For Figure 1E (Human ileum sections), the reference used was 100 nm above the most basal edge of the actin belt, determined by eye. Only enterocyte-to-enterocyte junctions were used in this study.

An approach similar to Figure 1B was used for Figure 2B,D (Caco-2 cells 6 and 9 days), but instead of the actin belt used as reference, we used the tight junction protein ZO-1, which signal was clearer.

#### Number of junctions used in the analyses

**Fig. 1B. Caco-2 cells 14 days**

Phall E-cad n= 13, Phall β-catenin n=11, Phall α-catenin n=10, Phall nectin-3 n=5, Phall afadin n=11, Phall ZO-1 n=12.

**Fig. 1E. Human ileum sections**

Phall E-cad, patient 1, n=7, patient 2, n=7, patient 3, n=5.
Phall β-catenin, patient 1, n=7, patient 2, n=7, patient 3, n=7.
Phall Nectin-2, patient 1, n=5, patient 2, n=8, patient 3, n=4.
Phall Afadin, patient 1, n=5, patient 2, n=10, patient 3, n=9.
Phall ZO-1, patient 1, n=9, patient 2, n=5, patient 3, n=8.

**Fig. 2B Caco-2 cells 6 days**

ZO-1 E-cad n=10, ZO-1 afadin n=9, Phall ZO-1 n=8.

**Fig. 2D Caco-2 cells 9days**

ZO-1 E-cad n=9, ZO-1 afadin n=9, Phall ZO-1 n=13.

#### Methods reference


1. I. Chantret, A. Rodolosse, A. Barbat, E. Dussaulx, E. Brot-Laroche, A. Zweibaum, M. Rousset, Differential expression of sucrase-isomaltase in clones isolated from early and late passages of the cell line Caco-2: evidence for glucose-dependent negative regulation. *J. Cell. Sci.* **107**, 213–225 (1994).

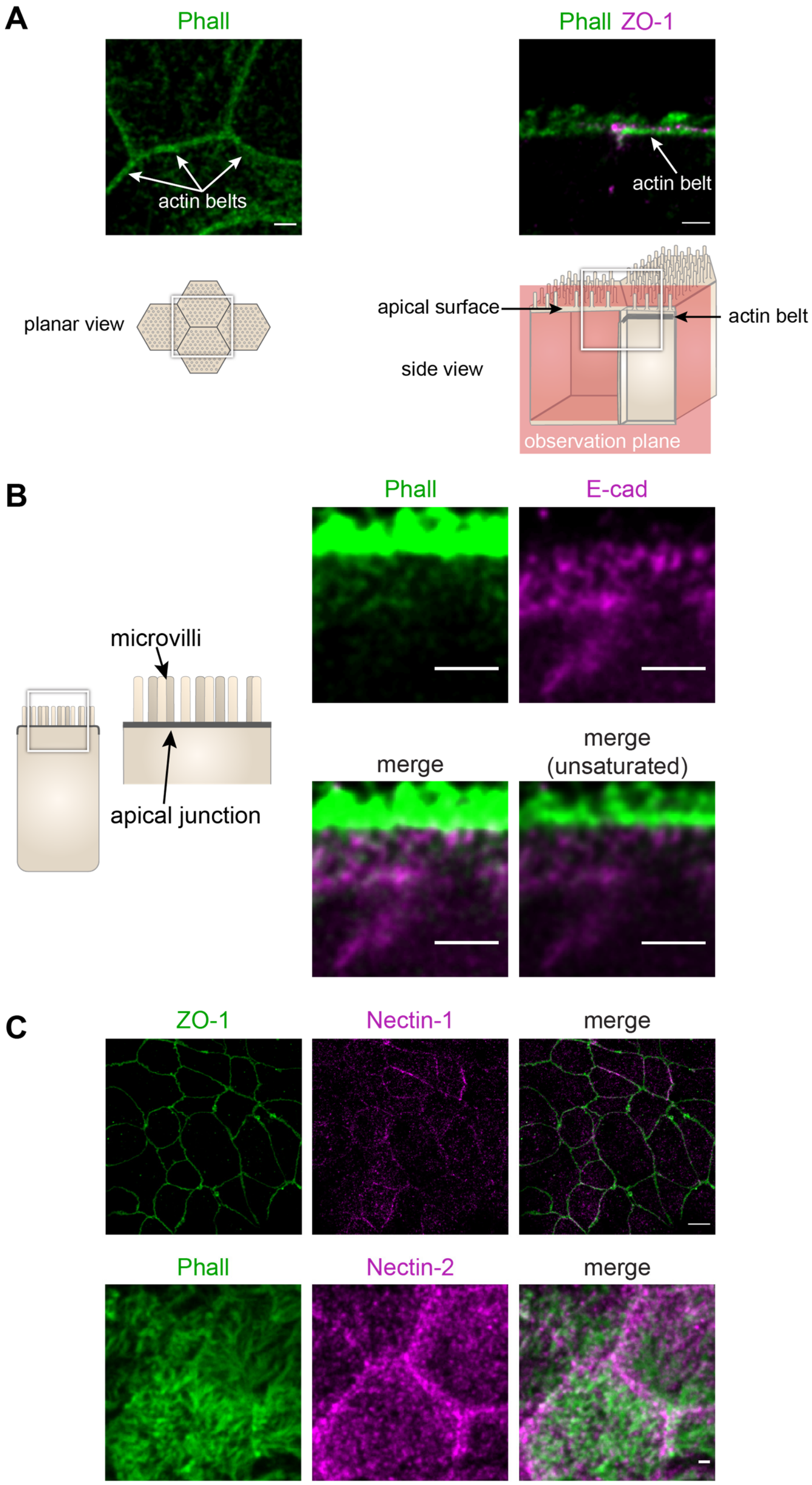


***Figure S1.*** Localization of the actin belt, E-cad, ZO-1, nectin-1 and nectin-2 in Caco-2 cells. (**A**) Actin belts are lining cell-to-cell apical junctions. Left, planar view, phalloidin staining. Right, side view, phalloidin (green), ZO-1 (magenta). Schematics represent the view of the image directly above. White squares represent the region where the image is taken. (**B**) E-cad is localized basally of the actin belt, where it colocalizes with dim F-actin structures. In the region of apical junctions, actin is present mostly in the actin belt and in microvilli. Side view, phalloidin (green), E-cad (magenta). (**C**) Nectin-1 and nectin-2 are localized at the apical junction. Planar views: top, ZO-1 (green), nectin-1 (magenta); bottom, phalloidin (green) labelling microvilli and the actin belt, nectin-2 (magenta). Scale bars: (A,B) 1 µm, (C) Top 5 µm, bottom 1 µm.

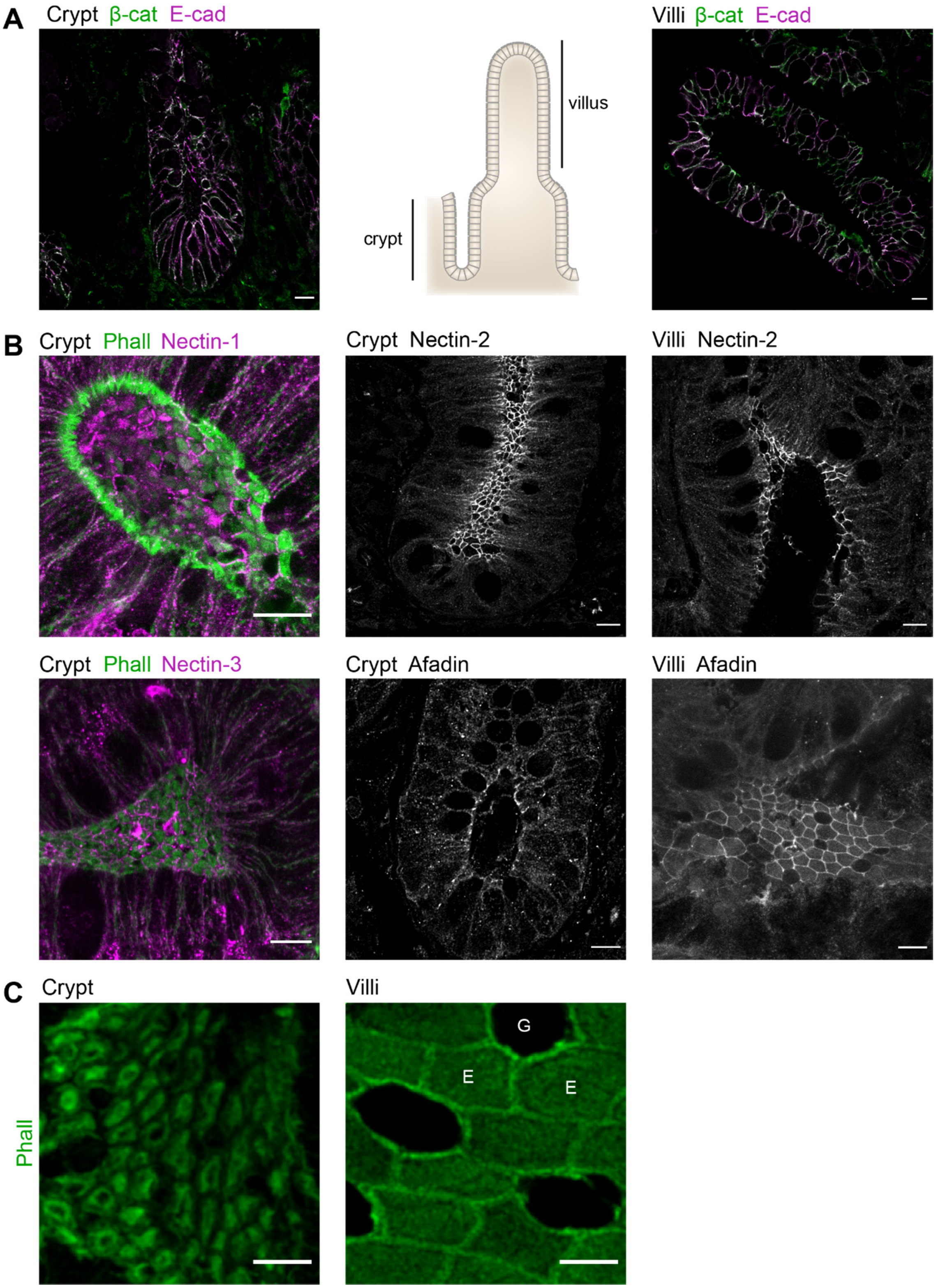


***Figure S2.*** Location of adhesive complexes and F-actin apical organization in human ileum biopsies. (**A**) E-cad and β-catenin in crypt (left) and villi (right). (Center) Schematic of villus and crypt regions at the surface of the small intestine. (**B**) Localization of nectin-1, nectin-2, nectin-3 and afadin in crypts and nectin-2 and afadin in villi. (**C**) F-actin organization at the apical surface of epithelial cells in villi (left) and crypt (right). "G" indicates a goblet cell with no apical F-actin, and "E" indicates two enterocytes. Phalloidin staining. Scale bars 10 µm.
